# Transcranial Brain-Wide Functional Ultrasound and Ultrasound Localization Microscopy in Mice using Multilinear Probes

**DOI:** 10.1101/2024.10.29.620796

**Authors:** Mathis Vert, Ge Zhang, Adrien Bertolo, Nathalie Ialy-Radio, Sophie Pezet, Bruno Osmanski, Thomas Deffieux, Mohamed Nouhoum, Mickael Tanter

## Abstract

Functional ultrasound imaging (fUS) and ultrasound localization microscopy (ULM) are advanced ultrasound imaging modalities for assessing both functional and anatomical characteristics of the brain. However, the application of these techniques at a whole-brain scale has been limited by technological challenges. While conventional linear acoustic probes provide a narrow 2D field of view and matrix probes lack sufficient sensitivity for 3D transcranial fUS, multilinear probes have been developed to combine high sensitivity to blood flow with fast 3D acquisitions. In this study, we present a novel approach the combined implementation of transcranial whole-brain fUS and ULM in mice using a motorized multilinear probe. This technique provides high-resolution, non-invasive imaging of neurovascular dynamics across the entire brain. Our findings reveal a significant correlation between absolute cerebral blood volume (ΔCBV) increases and microbubble velocity, indicating vessel-level dependency of the evoked response. However, the lack of correlation with relative CBV (rCBV) suggests that fUS cannot distinguish functional responses alterations across different arterial vascular compartments. This methodology holds promise for advancing our understanding of neurovascular coupling and could be applied in brain disease diagnostics and therapeutic monitoring.

## 1. Introduction

The recent advancements in functional ultrasound imaging (fUS) have opened new possibilities for utilizing ultrasound imaging in neuroscience research [1-5]. This technique leverages ultrafast Doppler imaging to estimate cerebral blood flow variations, thereby inferring the associated brain activity through neurovascular coupling [6, 7]. Its high resolution, single-trial sensitivity, and non-invasive nature make fUS an ideal ultrasound imaging modality for assessing brain function in preclinical animal models such as mice.

Complementarily, ultrasound localization microscopy (ULM) has introduced a super-resolution paradigm in ultrasound imaging, proving its effectiveness in mapping the microvasculature of various organs, including the brain and kidneys [8-10]. By using contrast-enhancing microbubbles (MBs) and ultrafast frame rates, ULM circumvents the diffraction-limited resolution barrier, revealing intricate details of hemodynamics even in deep seated organs [11, 12].

Despite the potential of these imaging modalities, several technical challenges remain, especially the transition from two-dimensional (2D) to three-dimensional (3D) imaging. To expand the field of view, two principle approaches have been proposed: motorized linear 1D probes [13, 14] and 2D matrix probes [15, 16]. However, the former lacks the fast-scanning capability necessary for whole-brain imaging at high frame rates, while the latter is often limited in sensitivity in imaging blood flow, particularly for transcranial brain imaging. Bertolo et al. recently introduced a motorized multilinear probe that combines the sensitivity of linear probes with the capability for fast whole-brain scanning [17].

Over the past decade, the concept of neurovascular coupling has significantly evolved to account for the cellular and structural diversity across various vascular compartments within the cellular vascular tree [18]. Literature suggests that the blood recruitment chronology goes from the smallest arterial compartments -capillaries and parenchymal arterioles-to the biggest -pial arterioles and cerebral arteries. While fUS can effectively estimate brain activity through neurovascular coupling, it remains unclear whether fUS alone can differentiate responses across different arterial levels, from pial arterioles and cerebral arteries to capillaries and parenchymal arterioles, for comprehensive functional analysis. Therefore, there is a need to perform fUS and ULM on the same imaging target and thus investigate whether functional activation in response to whisker stimulation correlates with changes in cerebral blood volume measured by fUS or microbubble velocity as estimated by ULM.

In this study, we built upon this work by demonstrating the feasibility of using such a probe to perform both fUS and ULM transcranially, enabling non-invasive whole-brain functional imaging in mice in conjunction with microvascular imaging and hemodynamic quantification at microscopic scale. We introduce an improved motorized scheme that optimizes imaging ratio (90% imaging ratio, with 10 seconds of imaging followed by 1.2 seconds of movement), facilitating enhanced MB detection within their relatively short lifetime. Additionally, we demonstrate that the resulting ULM maps encompass the whole-brain field-of-view, as achieved with fUS (11.9 × 7.87 × 8.1 mm^3^). We further investigate the registration of functional and anatomic data, allowing for detailed analysis of vascular structures in task-specific functional areas.

## 2. Methods

### 2.1 Ethics

All procedures were performed on six male C57BL/6J mice (7 weeks old, Janvier labs, France in compliance with European regulations and approved by the local ethics committee (Comité d’éthique en matière d’expérimentation animale number 59, Paris Centre et Sud, project number 25358 - 2020051019027581V2). The mice were housed three per cage with a 12-hour light/dark cycle, a constant temperature of 22°C and a humility level between 45 – 50 %. Unlimited access to food and water was provided. Before the experiments, animals were given a 1-week minimum acclimatization period to housing conditions. All experiments followed ARRIVE guidelines and relevant animal care regulations.

### 2.2 Animal Preparation

Anesthesia was induced with 2% isoflurane (delivered via a nose mask) and maintained using intraperitoneal medetomidine injection (80 µg/kg). The fur on the mice’s heads was shaved to prevent air bubbles within the ultrasound gel. Mice were fixed in a stereotaxic frame, and their eyes were protected with moisturizing gel (Ocry-Gel, Virbac, France). Body temperature was maintained at 37°C using a heating bed, monitored by a rectal probe. Heart rate and respiratory rate were monitored using a PowerLab data acquisition system with LabChart software (ADInstruments, USA). An intramuscular injection of medetomidine (100 µg/kg) will be performed after catheterization at the saphenous vein level for Microbubble injections (Sonovue, Bracco, Italy).Anesthesia levels were carefully adjusted to maintain stable physiological parameters, with isoflurane levels ranging from 0 to 0.5% and continuous medetomidine infusion at 100 µg/kg/h.

### 2.3 Multilinear Probe

Data was acquired transcranially using a multilinear ultrasound probe composed of four 64 elements linear sub-arrays with a center frequency of 15 MHz and a 0.11 mm pitch (IcoPrime-4D Multi-array, Iconeus, Paris, France). This probe was specifically designed for whole-brain fUS [17]. The probe was driven by a preclinical ultrasound system (Iconeus One, Iconeus, Paris, France) and mounted on a 4-axis motor system for rapid whole-brain scanning. The motorized setup moved in 525 µm steps— equivalent to one-quarter of the sub-array distance—enabling coverage of the whole brain with each step, as illustrated in Figure 1c. The imaging depth was approximately 8 mm, with a slice thickness of 500 µm.

**Figure 1.**
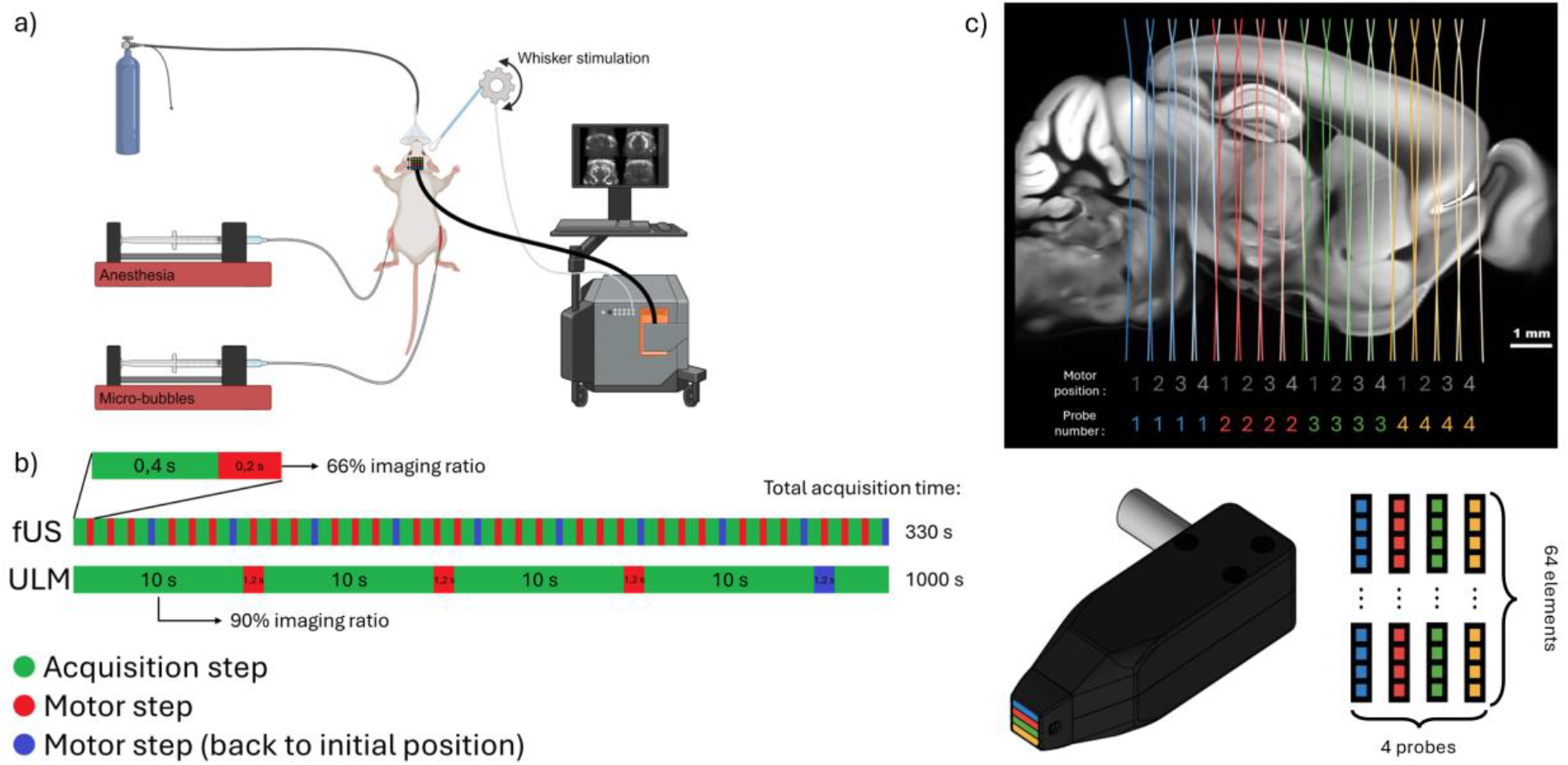
a) Setup schematic. Images were acquired using a preclinical ultrasound scanner. b) The proposed motor pattern enables 90% acquisition ratio to optimize the microbubble detection. c) The combination of a multilinear probe and a motorized setup enables fast whole brain scanning. The multilinear probe is composed of 4 linear 64 elements sub probes (15 MHz).

### 2.4 Imaging Acquisition and Beamforming

Functional data was captured using the scanner’s live acquisition software (IcoScan, Iconeus, Paris, France). ULM data was acquired using a custom version of the software. All data was acquired using tilted plane-wave imaging sequence with 10 angles compounded at 500 Hz. Beamforming was performed on a trapezoidal shape to enlarge the field of view using the maximum tilt angle of ± 12°.

### 2.5 Whisker Stimulation and Evoked Response Acquisitions

For fUS acquisitions, a mechanical index of 0.1 was used to enhance sensitivity and counteract signal attenuation due to the skull. The motor was programmed for 0.4-second acquisition periods, followed by 0.2-second movement intervals, resulting in a volume acquisition time of 2.4 seconds (0.42 Hz volume rate). Data from the four sub-arrays were concatenated for whole-brain analysis. During fUS acquisitions, mice whiskers were stimulated at 3 Hz using a cotton bud mounted on a servomotor controlled by an Arduino board, synchronized with the ultrasound scanner. Two stimulation protocols were used: 30 seconds baseline, 30 seconds ON, 30 seconds OFF (repeated five times), and 20 seconds baseline, 10 seconds ON, 20 seconds OFF (repeated ten times). Data from two animals subjected to the 10 seconds ON / 20 seconds OFF protocol were excluded due to artifacts.

### 2.6 Ultrasound Localization Microscopy Processing

After fUS acquisitions, ULM imaging was performed to achieve microscopic-resolution angiography. Sonovue© microbubble contrast agents (Bracco, Italy) were injected at a rate of 1.5 mL/h through the great saphenous vein. To optimize acquisition time, a customized motor protocol (Fig 1b) was implemented, achieving a 90% imaging ratio (10s acquisition, 1.2s motor movement). Data was collected over 1000 seconds, yielding 220 seconds of cumulative imaging per plane. Microbubble tracks were processed through a filtering-localization-tracking pipeline, achieving an in-plane resolution of 5 × 5 µm [8, 19].

Fourier ring correlation analysis was applied on ULM image to quantify the resolution limit of the final ULM image [20]. Briefly, all of the localization events were split into two subsets, and thus two sub-ULM images. The correlation of the spatial frequency content is then calculated as the normalized correlation of the two spectrum along iso-spatial frequency ring.

### 2.7 General Linear Model for Whisker Stimulation

Cerebral blood volume (CBV) changes were evaluated using a general linear model (GLM) to compare functional responses to expected stimuli patterns. Power Doppler signals were adjusted to eliminate baseline activity and enable the quantitative estimation of relative CBV (rCBV). Expected stimulus responses were modeled by convolving the stimulation pattern with a hemodynamic response function (HRF) selected from the literature [21]. The GLM design matrix included the stimulus and a constant slope to model signal drift. Statistical significance was determined using a t-test, and results were converted to z-scores for inter-subject comparison. P-values were corrected using the Bonferroni method.

### 2.8 Data Registration on Brain Functional Atlas

A brain position system (BPS) was leveraged to align the acquisitions to the Allen atlas before quantification [22]. Basically, a 3D Power Doppler volume is acquired at the beginning of each imaging session and then registered to a reference vascular map previously aligned with the brain atlas. This enables subsequent fUS acquisition of the same imaging session to be automatically aligned with the brain atlas for brain regions segmentation. As the fUS and ULM acquisitions were performed consecutively to be aligned during the image session, the same registration process can be used for ULM acquisitions. However, as the BPS system is performed using Power Doppler, the registration process average accuracy is approximately 100 μm. Thus, we performed additional manual registration to improve ULM acquisition alignment to the atlas.

### 2.9 Group-Level Statistics

For each fUS acquisition (n=10), quantitative rCBV and associated t-statistic were correlated with underlying MB velocity. The obtained correlation coefficients were redressed using the Fisher transform to enable group averaging. The metrics (rCBV and t-statistic) were tested against the hypothesis of a zero-mean using a Wilcoxon signed-rank test. The Wilcoxon signed-rank test is non-parametric and it does not assume a normal distribution of data. Therefore, it is suitable for providing robustness in assessing central tendencies of small sample sizes without strict distributional assumptions. Additionally, as it is a paired test, it focuses on the differences between paired observations, thus making it a sensitive test for detecting differences between two pairs of samples.

## 3. Results

### 3.1 Multilinear Probe Enables Whole-Brain Microscopic Angiography with ULM

The time-averaged Doppler signal from fUS and the rasterized MB counts from ULM are shown in Figure 2 for comparison. These images, acquired in two different modes, illustrate how the multilinear probe can scan the whole brain across 16 imaging planes. Both imaging modalities were performed during a single acquisition session using the same setup (probe, motors, scanner, and anesthesia), allowing the assessment of brain function using fUS and detailed microvascular anatomy using ULM. Doppler images demonstrate high sensitivity to blood flow, except in the first three frontal slices where the skull absorption and aberrations impede transcranial imaging. Doppler sensitivity plays a critical role in functional analysis by inferring neuronal activation based on variations in CBV signal over time. The Doppler images result from averaging signals over a task-based acquisition lasting 330s, as can be seen from Figure 1b. ULM localization map enables high-resolution mapping of microvasculature, resolving vessels as small as 32 µm, as can be seen from Figure 2d. They are the result of the rasterization of 2.5 million bubbles tracked in 1000 s. The local comparison of Doppler and ULM maps in Figure 2c shows how ULM reveals vessels hidden in Doppler maps, much smaller than the pixel size. In Figure 2c the blue arrows point to regions where the vessels do not raise contrast in Doppler images while being well defined in ULM. The intensity profile along the green line was extracted from both maps and compared in Figure 2d.

**Figure 2.**
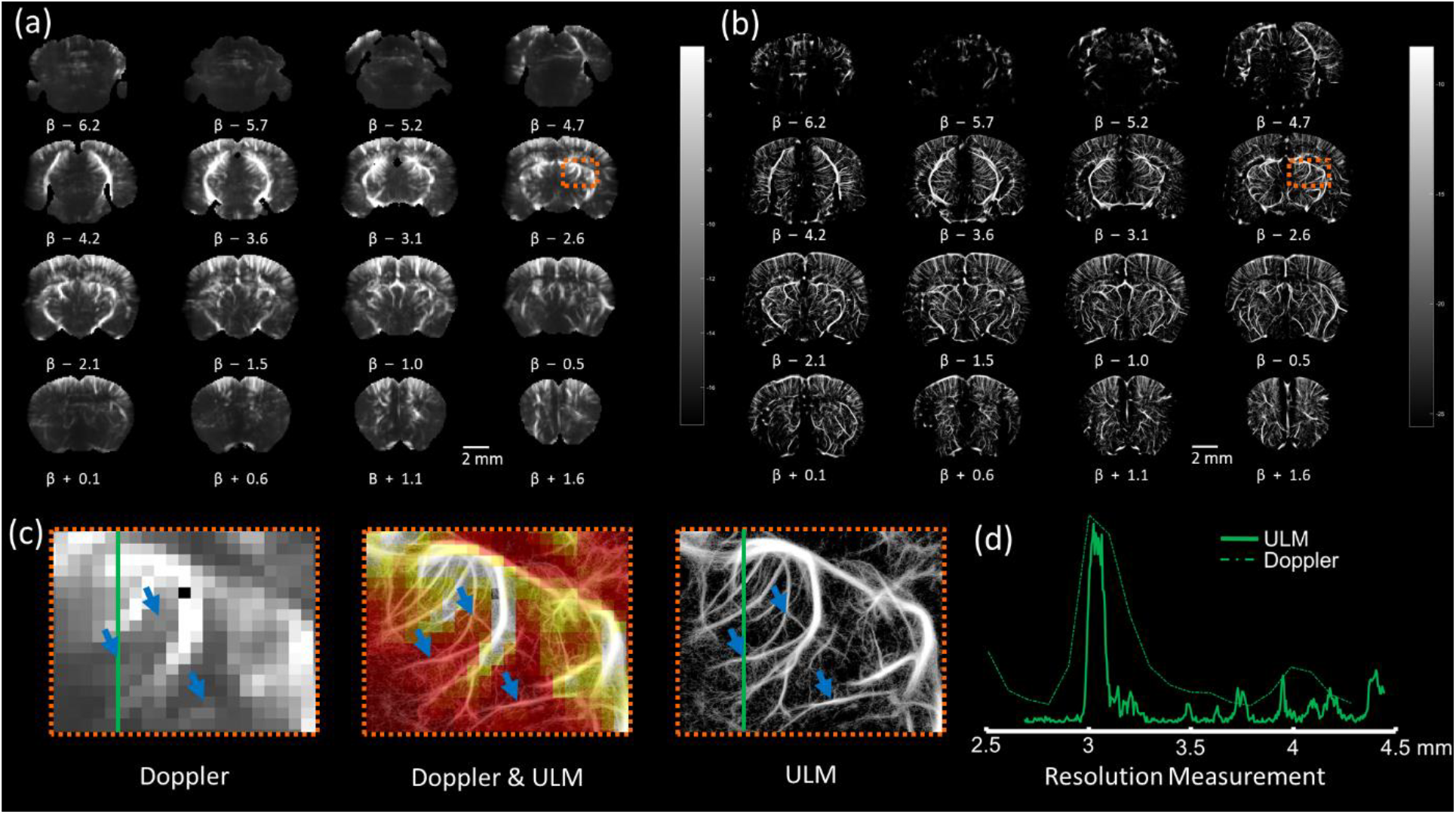
Comparison between Doppler image and ULM localization maps, covering a field of view of 11.9 × 7.87 × 8.1 mm^3^ across 16 imaging planes. (a) Time-averaged Doppler images highlight major vascular structures at a resolution of 110 × 98 × 525 µm. (b) Microbubble localization maps provide a more detailed representation of microvascular features at 5 × 5 × 525 µm resolution. (c) Overlaid Doppler and ULM images demonstrate how ULM captures finer microvascular structures, as indicated by blue arrows. These arrows mark regions where ULM reveals microvasculature which cannot be visualized in Doppler images. (d) The intensity profile, drawn along a green line, contrasts the resolution limits of Doppler and ULM, with Doppler resolving vessels down to 234 µm and ULM detecting structures as small as 32 µm.

### 3.2 Whole-Brain Quantification of Blood Flow Properties via MB Tracking

In Figure 3a, time-averaged Doppler images effectively depict major vascular structures, achieving a spatial resolution of 110 × 98 × 525 µm. This resolution captures primary vasculature but lacks the details needed to resolve finer microvascular networks. By contrast, Figure 3b showcases microbubble velocity maps obtained through ULM, revealing a highly detailed representation of microvascular flow information with a significantly enhanced resolution of 5 × 5 × 525 µm. This improvement enables visualization of microvascular flow information that Doppler imaging cannot distinguish. In Figure 3c, the combined Doppler and ULM images provide complementary insights, highlighting additional vascular structures detected by ULM, thus enhancing the depth of structural detail across the vascular network. Lastly, Figure 3d employs Fourier ring correlation analysis on the ULM image, determining a resolution limit of 13.33 µm. This analysis substantiates the superior resolving power of ULM for microvascular imaging, achieving a level of detail that Doppler imaging alone cannot provide.

**Figure 3.**
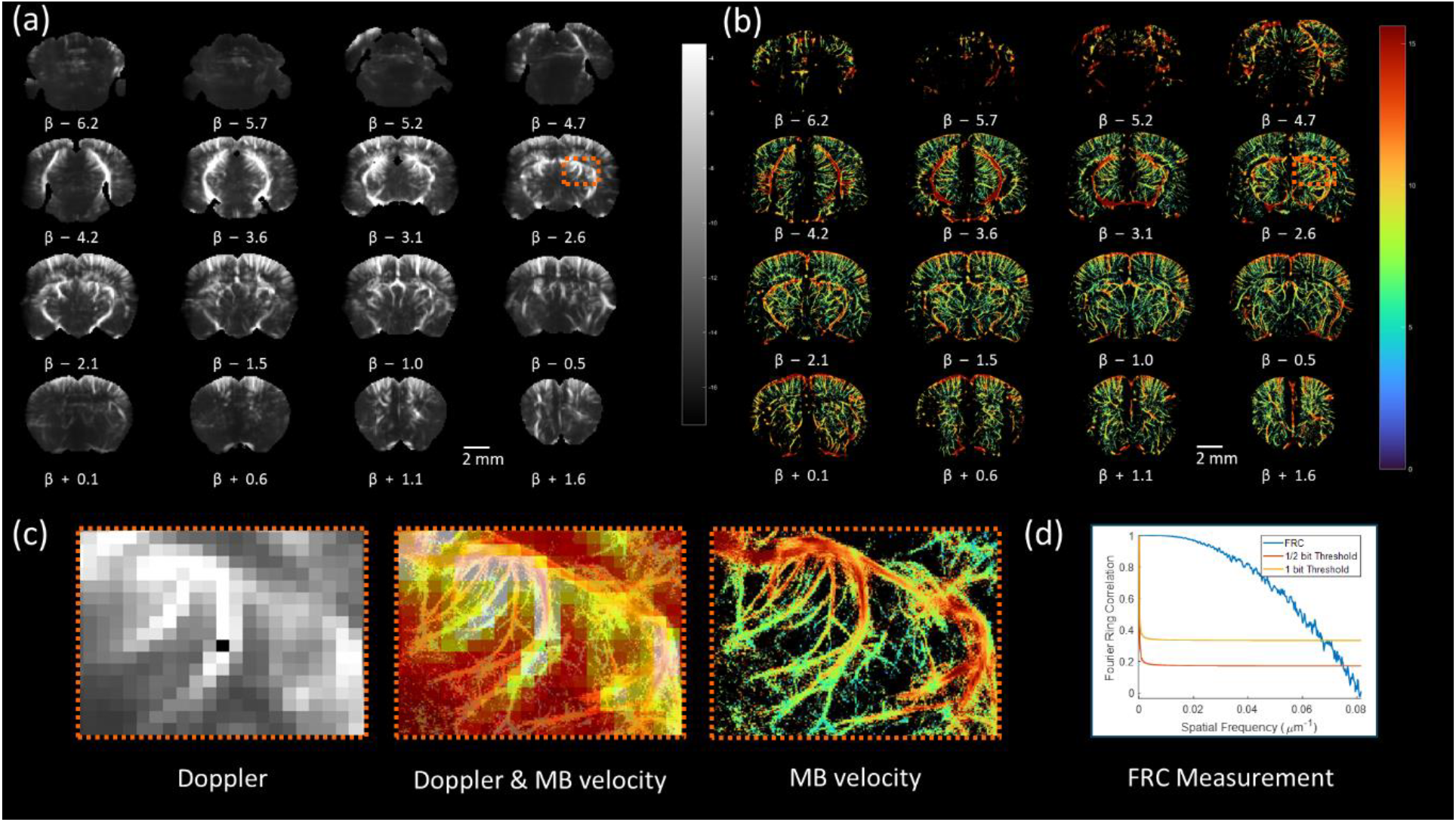
Comparison between Doppler image and ULM velocity maps, covering a field of view of 11.9 × 7.87 × 8.1 mm3 across 16 imaging planes. (a) Time-averaged Doppler images highlight major vascular structures at a resolution of 110 × 98 × 525 µm. (b) Microbubble velocity maps provide a more detailed representation of microvascular features and blood flow velocity at 5 × 5 × 525 µm resolution. (c) The combined Doppler and ULM images reveal additional vascular structures captured by ULM. (d) Fourier ring correlation analysis on ULM image, demonstrates a resolution limit of 13.33 µm.

### 3.3 ULM Reveals Microvasculature in Functionally Active Regions

Figure 4 summarizes the results of the assessment of the brain function through whisker stimulation and the underlying network of micro vessels. The maps shown in purple display the z-score of the pixel-wise correlation between the task stimulus and the measured Doppler signal. The measured response to the stimuli is significant for single trial fUS acquisition. The significantly activated area fits the expected functional region, i.e. the barrel field region of the cortex (as can be seen from Figure 4a, b). Figure 4b shows how the results are reproducible for each animal. Also, the spatially averaged time signal of the barrel field is highly correlated with the temporal track of the stimulus (Figure 4c). The measured relative increase of CBV in this region reaches 15% during the stimulation.

**Figure 4.**
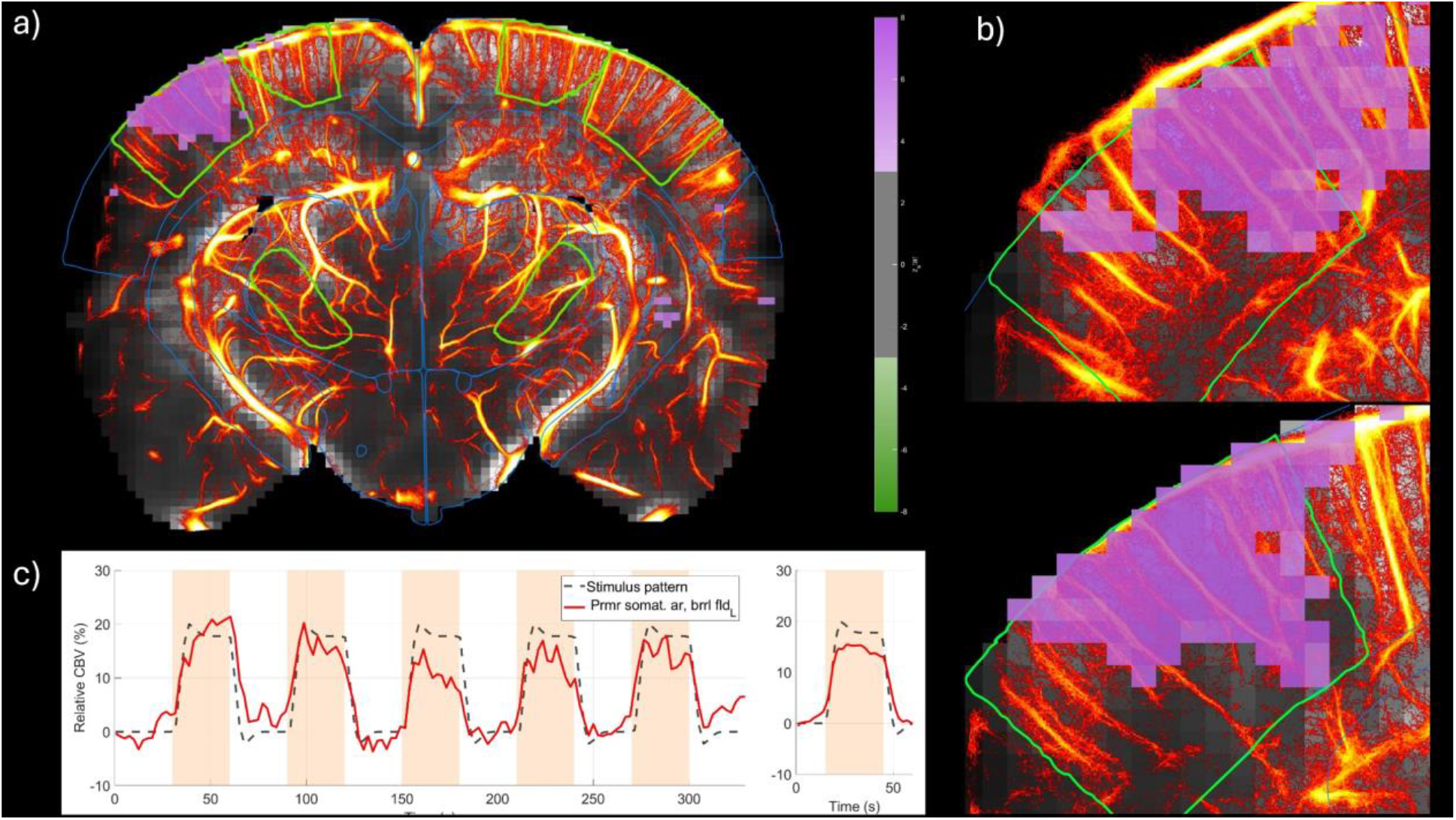
Demonstration of ULM exposes the microvasculature underlying functional brain regions, in response to whisker stimulation. (30-second ON, 30-second OFF). (a) A brain slice intersecting the barrel cortex, with a z-score map highlighting the contralateral cortical response to whisker stimulation. (b) Zoom-in images of functionally active areas from two animals demonstrate the reproducibility of the stimulation and its mapping to the expected region. (c) Temporal analysis shows strong correlation between the barrel cortex signal and the stimulation pattern, with a 15% relative increase in CBV during whisker stimulation.

### 3.4 Group-Level Statistics Reveal the Hierarchy of Functional Responses Across Vessels

The neurovascular response to a stimulus is supposedly varying depending on the arterial level. As shown in Figure 5a, ULM velocity maps can describe such a hierarchy of arterial vessels while Doppler and thus fUS images do not. However, while fUS does not have the spatial resolution to distinguish this range of vessels, it still is sensitive to slow flows down to 1mm/s (Figure 5b). We can interpret this as a spatial averaging of the signal coming from arterioles level 1 to 4 (see green rectangle) in 100×100×525µm^3^ voxels. As demonstrated in Figure 5c, over the cohort (n=10), the absolute increase of CBV (ΔCBV) is significantly correlated with the MB velocity (r = 0.19, p =1.9*10-3). This means that the strength of the evoked response is correlated with the vessels level. On contrary, the relative response – measured as the relative increase of CBV-is on average not correlated with the velocity.

**Figure 5.**
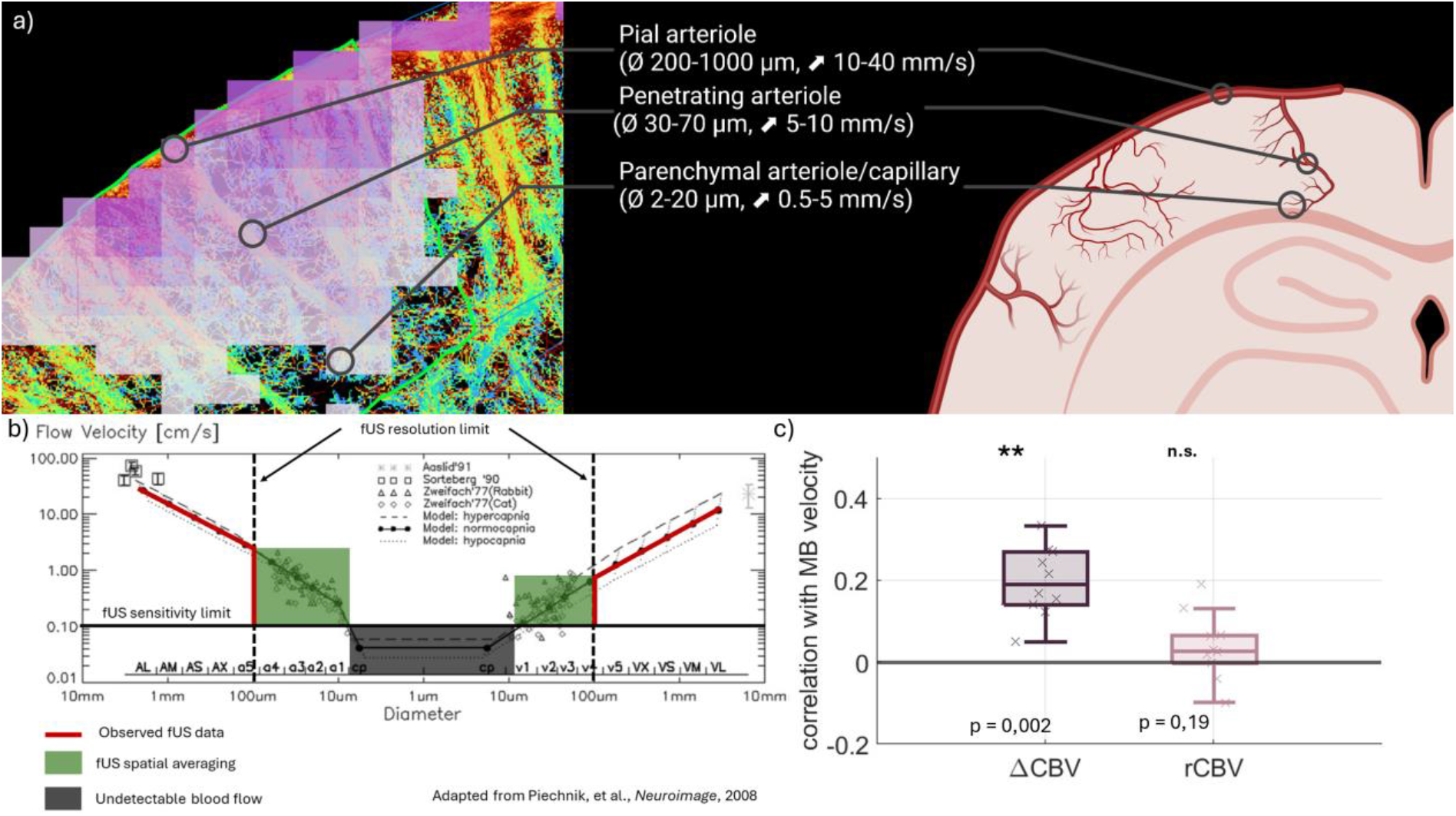
Relationship between vessels rank and functional response, using MB velocity as a proxy for vessel hierarchy. (a) Overlay of the relative CBV increase in the barrel field and the MB velocity, delineating different arterial compartments based on diameter and velocity. (b) fUS sensitivity and resolution thresholds draw the limits of detection of the vascular tree based on its spatial features (flow velocity and diameter dependence corresponds to Fig. 5b from Piechnik et al, Neuroimage, 2008). The solid black line defines the slowest flows that fUS is sensitive to. Dotted black line defines the voxel size in which any sources of signal are averaged (green area). (c) Distribution of the correlation of ΔCBV and rCBV with MB velocity. The absolute evoked response (ΔCBV) is correlated with measured velocity. The response significance (rCBV) is decorrelated from the flow due to the lack of fUS resolution on the lowest arterial branches. (*: p<0.05, **: p<0.01, n =10 acquisitions, 6 mice)

## 4. Discussion

This study demonstrated the successful application of functional ultrasound imaging (fUS) and ultrasound localization microscopy (ULM) using a motorized multilinear probe to achieve transcranial whole-brain functional and microvascular imaging in mice. By combining these two modalities in a single setup taking benefit of the high sensibility of the multilinear probe technology, it was able to capture both functional and anatomical data, and introduce a tool that may offer comprehensive insights into the cerebral microvasculature and its role in neurovascular coupling. The optimized motorized scanning scheme allowed for a 90% imaging ratio, enhancing microbubble (MB) detection, while the ULM maps provided detailed microvascular structures not visible in power Doppler images. Our results indicate that functional activation in response to whisker stimulation was strongly correlated with changes in cerebral blood volume (CBV), and that this response is not dependent on MB velocity. This suggests that Doppler-based filtering of low velocities may not be sufficient to fully capture the complexity of the neurovascular response in particular the discrepancy of relative neurovascular response in arterioles and smaller capillary vessels.

As can be seen from Figure 5, the vessels hierarchy from ULM maps and the observed spatial distribution of vessels and functional response (shown in red line) can be inferred given by fUS. In other words, if we make the strong assumption that small velocities correspond to small vessels, it could potentially be possible to distinguish the functional response of small vessels from the one of larger arterioles by separating these components in the Doppler Fourier spectrum. To evaluate if such inference was possible from the data, we measured the correlation between the fUS functional response and the underlying microvessels blood velocity. We considered the MB velocity as a proxy of the blood velocity and consequently of the vessel hierarchical level. The results demonstrated in Figure 5c suggest that this result denies the possibility to distinguish the response of different arterial levels in the Doppler data. Indeed, both Two-Photon optical imaging and functional ULM have shown that the relative CBV increase during stimulation should be much higher in the Arteriolar-Capillary Transition (ACT) zone than in the penetrating arterioles [23-26]. We clearly see in Figure 4b that the rCBV change is not higher in the regions located in between two penetrating arterioles compared to the rCBV change in pixels located in a penetrating arteriole.

Neurovascular coupling, the process by which neuronal activity is linked to changes in cerebral blood flow, requires detailed knowledge of both vascular structure and flow dynamics. Conventional techniques such as fMRI provide only macroscopic views of this process, whereas fUS allows for high-resolution, real-time measurement of blood flow changes in response to neuronal activation [27]. ULM, with its ability to track microbubbles and reveal capillary-level details, adds a new dimension by mapping the microvascular network responsible for these changes. By correlating functional activation data with microvascular structure and blood velocity, this study enhances our understanding of how different vascular compartments contribute to neurovascular coupling. The ability to distinguish between different vessel sizes and flow rates also reveals how blood is recruited during neuronal activation, providing a more comprehensive view of the underlying hemodynamic processes.

The non-invasive nature of fUS and ULM, combined with their ability to provide high-resolution microvascular maps, makes them ideal tools for longitudinal studies. These techniques allow for repeated imaging of the same animal over time, enabling researchers to monitor changes in the microvasculature under different conditions. This could be particularly useful for studying microvascular alterations in neurodegenerative diseases, stroke recovery, or chronic conditions such as diabetes [28]. Monitoring changes in vessel density, flow velocity, and vascular dynamics over time would provide valuable insights into disease progression and the effects of therapeutic interventions. Longitudinal studies using fUS and ULM could also be applied to track vascular plasticity in response to environmental or behavioral changes, further deepening our understanding of brain health and resilience.

The ability to differentiate between various vascular and neuronal pathologies is one of the most promising clinical applications of fUS and ULM. Vascular dementia and Alzheimer’s disease, for example, share similar symptoms but are driven by different underlying pathologies. Vascular dementia is primarily caused by reduced blood flow to the brain, often due to microvascular damage, while Alzheimer’s is characterized by amyloid-beta accumulation and neurodegeneration [29]. ULM’s ability to map microvascular structures with high resolution could be instrumental in distinguishing these conditions. By quantifying changes in microvascular density and flow patterns, ULM and fUS could help identify specific vascular abnormalities that are characteristic of vascular dementia, such as microinfarcts or capillary loss. This would allow for earlier and more accurate diagnosis, enabling better-targeted interventions for each condition [30].

Vascular plasticity refers to the brain’s ability to remodel its vascular network in response to changes in neuronal activity, metabolic demand, or injury. This plasticity is critical for maintaining proper brain function, particularly in aging and disease [31]. The ability of ULM to visualize and quantify microvascular dynamics at high resolution provides a unique opportunity to study this process. By tracking changes in vessel diameter, flow velocity, and microvascular density over time, ULM can reveal how the brain adapts to new demands or compensates for damaged regions. Furthermore, longitudinal studies using fUS and ULM could investigate how vascular plasticity is affected by various factors, such as aging, exercise, or pharmacological treatments, providing new insights into therapeutic strategies for maintaining brain health [32].

## 5. Conclusion

This study successfully demonstrates the feasibility of performing whole-brain fUS and ULM using a motorized multilinear probe in mice. By integrating these two advanced imaging techniques, we were able to capture both functional and anatomical data, offering a comprehensive view of the neurovascular dynamics in the mouse brain. The motorized scanning scheme enabled a high imaging ratio, optimizing microbubble detection and yielding high-resolution ULM maps that revealed intricate microvascular details beyond the scope of Doppler images. Furthermore, the study highlights that the functional response to stimuli, as assessed by CBV changes, is not directly correlated with microbubble velocity, suggesting that velocity-based Doppler filtering alone may not fully capture the complexity of neurovascular coupling. This combined approach provides a more detailed understanding of neurovascular coupling by revealing the hierarchical recruitment of blood flow across different vascular compartments, paving the way for future applications in both preclinical and clinical neuroscience. Additionally, this methodology could be applied in longitudinal studies to monitor microvascular alterations in neurodegenerative diseases or assess vascular plasticity. The ability to distinguish functional vascular changes could also facilitate the diagnosis of vasculo-neuronal pathologies, such as vascular dementia and Alzheimer’s disease, further enhancing the potential of this imaging modality for clinical translation.

## Acknowledgement

This work was supported by Inserm research accelerator (Inserm ART) in Biomedical Ultrasound and by the French national research agency (ANR) under ANR-21-CE19-0050 program (Project SonoGT). This research was also funded by the Region Ile de France - Convention DIM ELICIT Innovative Technologies for Life Science.

